# Mechanistic analysis of a novel membrane-interacting variable loop in the pleckstrin-homology domain critical for dynamin function

**DOI:** 10.1101/2022.09.03.506501

**Authors:** Himani Khurana, Soumya Bhattacharyya, Thomas Pucadyil

## Abstract

Classical dynamins are best understood for their ability to generate vesicles by membrane fission. During clathrin-mediated endocytosis (CME), dynamin is recruited to the membrane through multivalent protein and lipid interactions between its Proline Rich Domain (PRD) with SRC Homology 3 (SH3) domains in endocytic proteins and its Pleckstrin Homology Domain (PHD) with membrane lipids. Variable loops (VL) in the PHD bind lipids and partially insert into the membrane thereby anchoring the PHD to the membrane. Recent molecular dynamics (MD) simulations reveal a novel VL4 that interacts with the membrane. Importantly, a missense mutation that reduces VL4 hydrophobicity is linked to an autosomal dominant form of Charcot Marie Tooth (CMT) neuropathy. To mechanistically link data from simulations with the CMT neuropathy, we analyzed functions of a VL4 mutant with reduced hydrophobicity. In assays that rely solely on lipid-based membrane recruitment, this mutant showed an acute membrane curvature-dependent binding and fission. Remarkably, in assays that mimic a physiological multivalent lipid- and protein-based recruitment, this mutant was completely defective in fission across a range of membrane curvatures. Importantly, expression of this mutant in cells inhibited CME, consistent with the autosomal dominant phenotype associated with the CMT neuropathy. Together, our results emphasize the significance of finely tuned lipid and protein interactions for efficient dynamin function.

## Introduction

Dynamins contain a GTPase (G) domain, a stalk domain and a bundle-signaling element (BSE). The G domain hydrolyzes GTP, and the stalk maintains dynamin as a tetramer in solution. The stalk also facilitates self-assembly into helical scaffolds on the membrane. The BSE transmits self-assembly induced conformational changes in the scaffold to the G domain causing stimulation in dynamin’s basal GTPase activity (Faelber et al., 2012; Chappie and Dyda, 2013; Schmid and Frolov, 2011). In addition to these core domains, classical dynamins contain an additional membrane binding Pleckstrin Homology Domain (PHD) and a Proline Rich Domain (PRD). The PHD binds phosphatidylinositol-4,5-bisphosphate (PI(4,5)P_2_) and phosphatidylserine (PS) while the PRD binds SRC Homology 3 (SH3) domains in endocytic accessory proteins. Multivalent binding to endocytic proteins and lipids serves to recruit dynamin to emergent membrane buds during clathrin-mediated endocytosis (CME) (Ferguson and Camilli, 2012; Antonny et al., 2016; Mettlen et al., 2009; Daumke et al., 2014).

The PHD is a hotspot for mutations that are linked to several inherited genetic disorders. Mutations causing centronuclear myopathy (CNM) and Charcot-Marie-Tooth (CMT) neuropathy are predominantly located in this domain (Durieux et al., 2010; Haberlová et al., 2011; Fabrizi et al., 2007; Bitoun et al., 2008; Zuchner et al., 2005; Kenniston and Lemmon, 2010). CNM mutations map to a region on the PHD distant from those required for lipid binding while CMT mutations are frequently localized in the PHD and affect lipid binding (Kenniston and Lemmon, 2010). The crystal structure of the PHD reveals a core β-sandwich that is formed of two sheets oriented in an antiparallel arrangement and a single α-helix (Ferguson et al., 1994; Lemmon, 2000). The β strands are connected by variable loops (VLs). VL1 (^531^IGIMKGG) is a membrane-inserting loop and mutations that reduce its hydrophobicity render dynamin defective in membrane binding, fission, and CME (Ramachandran et al., 2009; Dar et al., 2015; Shnyrova et al., 2013). Recent molecular dynamics (MD) simulations of the isolated PHD confirm membrane insertion of the VL1, with the terminal atom on the I533 residue at the tip of VL1 dipping 0.58 nm below the phosphate plane (Baratam et al., 2021). The relatively polar VL2 (^554^KDDEEKE) and VL3 (^590^NTEQRNVYKDY) do not directly interact with the membrane (Baratam et al., 2021), but are important for dynamin functions (Züchner et al., 2005) (Liu et al., 2011).

The PHD structure shows a fourth VL (VL4) (^576^EKGFMSSK) located between β5 and β6 strands and MD simulations show that it interacts directly with the membrane (Baratam et al., 2021). This is surprising considering that VL4 has much lower hydrophobicity than VL1. VL4 therefore represents a novel membrane interacting loop which together with VL1 anchors the PHD on to the membrane. Importantly, a missense mutation M580T in VL4 has been linked to an autosomal dominant form of CMT (Haberlová et al., 2011). MD simulations reveal that the adjacent F579 residue in VL4 approaches closest to the membrane, with the terminal atom dipping 0.41 nm below the phosphate plane (Baratam et al., 2021). Surprisingly, simulations with the F579A mutant showed no apparent defect in membrane binding. Instead, this mutant, missing one of the two membrane interacting pivots, rendered the PHD to sample a wider range of rotational conformations about the membrane normal. To mechanistically link data from MD simulations to the CMT neuropathy, we analyzed the VL4 mutant (F579A) in a suite of in vitro assays analyzing membrane binding and fission and in cellular assays monitoring the effects of expression of this mutant on CME.

## Results

The VLs show a high degree of conservation across dynamin isoforms and across dynamins from different species (Supplementary Fig. 1). The PHD contains positively charged clusters towards the membrane binding interface (Fig. 1A). MD simulations reveal that lysine residues outside the VLs (K359, K554) and within VL1 (K535) recruit the anionic lipid DOPS towards the PHD and stabilize its membrane association (Baratam et al., 2021). To rule out potential global defects caused by the F579A mutation in VL4 and establish the validity of our binding and fission assays, we first analyzed dynamin functions on highly anionic DOPS-containing membranes. Membrane binding was tested using a Proximity-based Labeling of Membrane Associated Proteins (PLiMAP) assay (Jose et al., 2020; Jose and Pucadyil, 2020). In this assay, proteins are mixed with liposomes containing 1 mol% of a bifunctional fluorescent crosslinking lipid. Membrane binding brings the protein in proximity of the bifunctional lipid, which upon UV exposure becomes crosslinked with the lipid. Samples are then resolved using SDS-PAGE and binding can be analyzed by measuring lipid fluorescence associated with the protein. PLiMAP assays revealed strong binding of both Dyn1 and Dyn1(F579A) to DOPS liposomes (Fig. 1B). Furthermore, membrane binding caused robust stimulation in the GTPase activities of both Dyn1 and Dyn1(F579A) (Fig. 1C). On Supported Membrane Templates (SMrTs), which comprise of an array of membrane nanotubes supported on PEG cushions (Dar et al., 2015, 2017), the addition of Dyn1(F579A) with GTP caused fission and severing of nanotubes, just like Dyn1 (Fig. 1D). Together, these results rule out any global effects on protein function caused by introducing the F579A mutation in Dyn1.

**Fig. 1.**
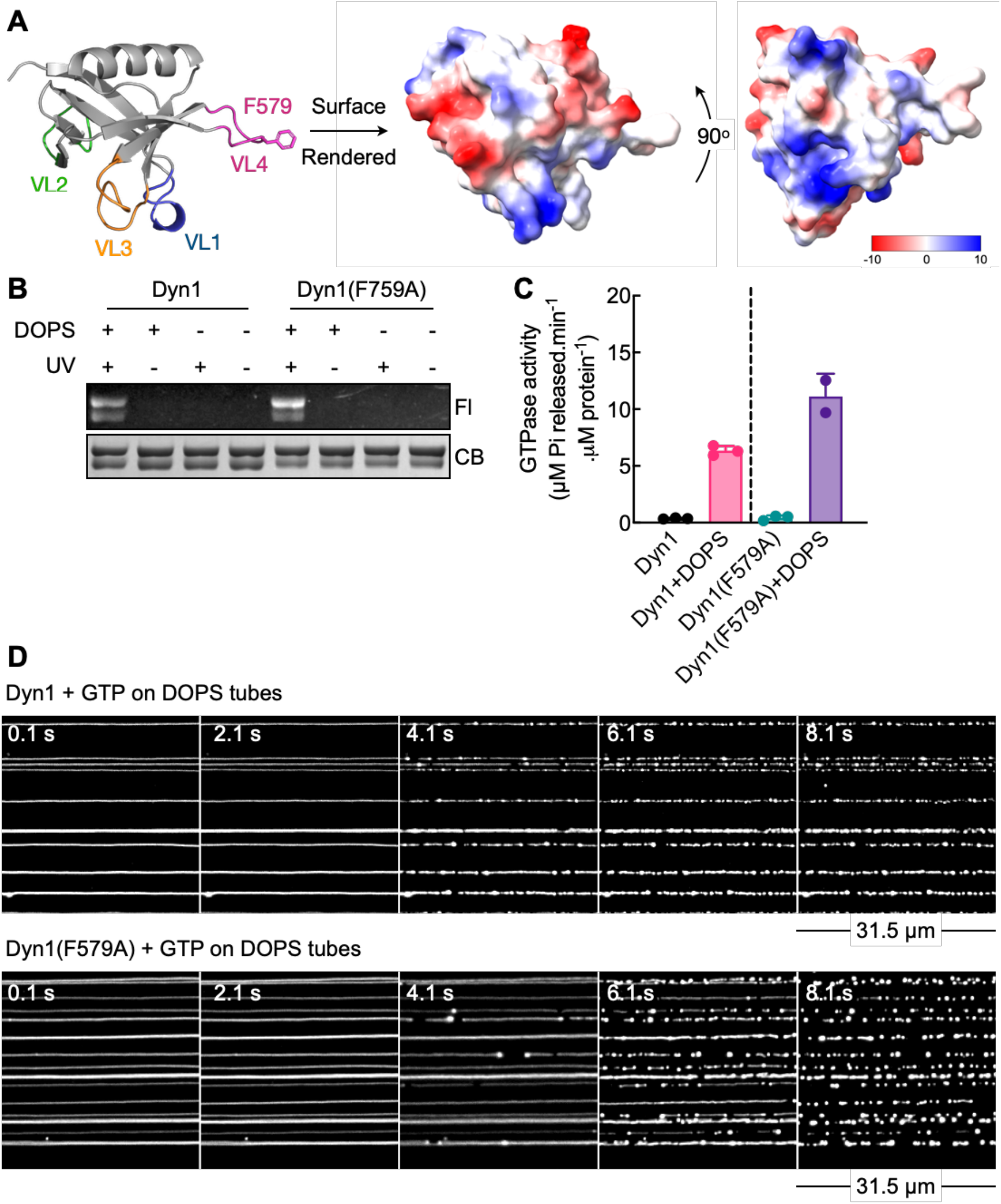
Analyzing the VL4 mutant on highly anionic membranes. (A) Crystal structure of the dynamin1 PHD (PDB: 1DYN), showing the 4 variable loops (VL1-4). The F579 residue in VL4 is marked. Also shown are space-filling models of the structure showing positively charged surface patches present on the membrane binding side. Models were rendered using ChimeraX (Jurrus et al., 2018; Pettersen et al., 2020). (B) Results from a representative PLiMAP experiment showing in-gel fluorescence (Fl) and Coomassie brilliant blue (CB) staining of Dyn1 and Dyn1(F579A) on DOPS liposomes. Dynamin migrates as a doublet with the low molecular weight band representing a partial truncation at the C-terminus (see Methods and (Dar and Pucadyil, 2017)). (C) GTPase activity of Dyn1 and Dyn1(F579A) in the absence and presence of DOPS liposomes. Data represent the mean ± SD of three independent experiments. (D) Frames from a time-lapse movie showing fission of DOPS SMrTs with Dyn1 and Dyn1(F579A) in the presence of GTP.

Due to the sensitive fluorescence-based read-out, PLiMAP is particularly useful in testing binding of proteins to liposomes containing low and physiologically relevant concentrations of lipids. We tested binding of Dyn1 and Dyn1(F579A) to liposomes containing 1 mol% PIP_2_ and 15 mol% DOPS. Dyn1 bound these membranes with an apparent affinity (K_d_) of ∼200 nM, which is consistent with previous estimates (Kenniston and Lemmon, 2010). Under these stringent conditions, Dyn1(F579A) showed substantial defects with a significantly lower binding affinity (K_d_ ∼ 560 nM) and a 2.6-fold lower maximal binding (B_max_) (Fig. 2A,B). While a reduction in binding affinity of Dyn1(F579A) is consistent with the reported interaction between the VL4 with the membrane, a reduction in B_max_ was unexpected and we investigated the cause for this effect. Previous reports analyzing binding of peripheral membrane proteins have indicated that while the K_d_ reflects the strength of interaction between specific residues on the protein with lipids, the B_max_ depends on abundance of both the interacting lipid as well as membrane defects that facilitate insertion of hydrophobic residues in proteins (Hatzakis et al., 2009). Furthermore, membrane defects are more abundant on membranes of high curvature (Vanni et al., 2014). MD simulations reveal that unlike VL1, VL4 does not directly interact with PI(4,5)P_2_ (Baratam et al., 2021) and we wondered if the ability of VL4 to insert into membrane defects could allosterically stabilize dynamin on the membrane. If this was the case, then the F579A mutation could render dynamin binding more sensitive to membrane curvature. To test this, we turned to a spatially resolved microscopic assay that monitors protein binding to membrane nanotubes of different sizes (and hence curvatures) in SMrTs. We have earlier confirmed that PI(4,5)P_2_ is uniformly distributed on membrane nanotubes of varying sizes (Dar et al., 2015), which should allow us to specifically test the influence of membrane curvature on protein binding. SMrTs were incubated with fluorescent Dyn1 and Dyn1(F579A) for 15 min, washed with buffer and imaged. Dyn1 binding and self-assembly caused it to organize as numerous foci on every nanotube (Fig. 2C). In contrast, Dyn1(F579A) formed substantially fewer foci, which were only found on some tubes (Fig. 2C). To analyze the dependence of membrane curvature on dynamin binding, we correlated the probability of detecting a dynamin focus on a nanotube to its radius. For Dyn1, such binding probability showed a shallow dependence on the starting tube size (Fig. 2D). In contrast, Dyn1(F579A) was severely defective with significantly lower binding seen even on highly curved tubes and a sharp decline in binding on wider tubes (Fig. 2D). Dyn1 foci on SMrTs are scaffolds that are capable of constricting and severing tubes in presence of GTP (Dar et al., 2015; Dar and Pucadyil, 2017). Consistent with the trend seen with membrane binding (Fig. 2D), addition of Dyn1 with GTP showed robust fission of nanotubes of a wide range of sizes while the addition of Dyn1(F579A) showed severe defects in fission, with tubes thinner than 15 nm radius showing a fractional fission probability and tubes wider than 20 nm radius showing no fission (Fig. 2E). Thus, compromising the hydrophobicity of VL4 causes dynamin functions to become acutely sensitive to membrane curvature.

**Fig. 2.**
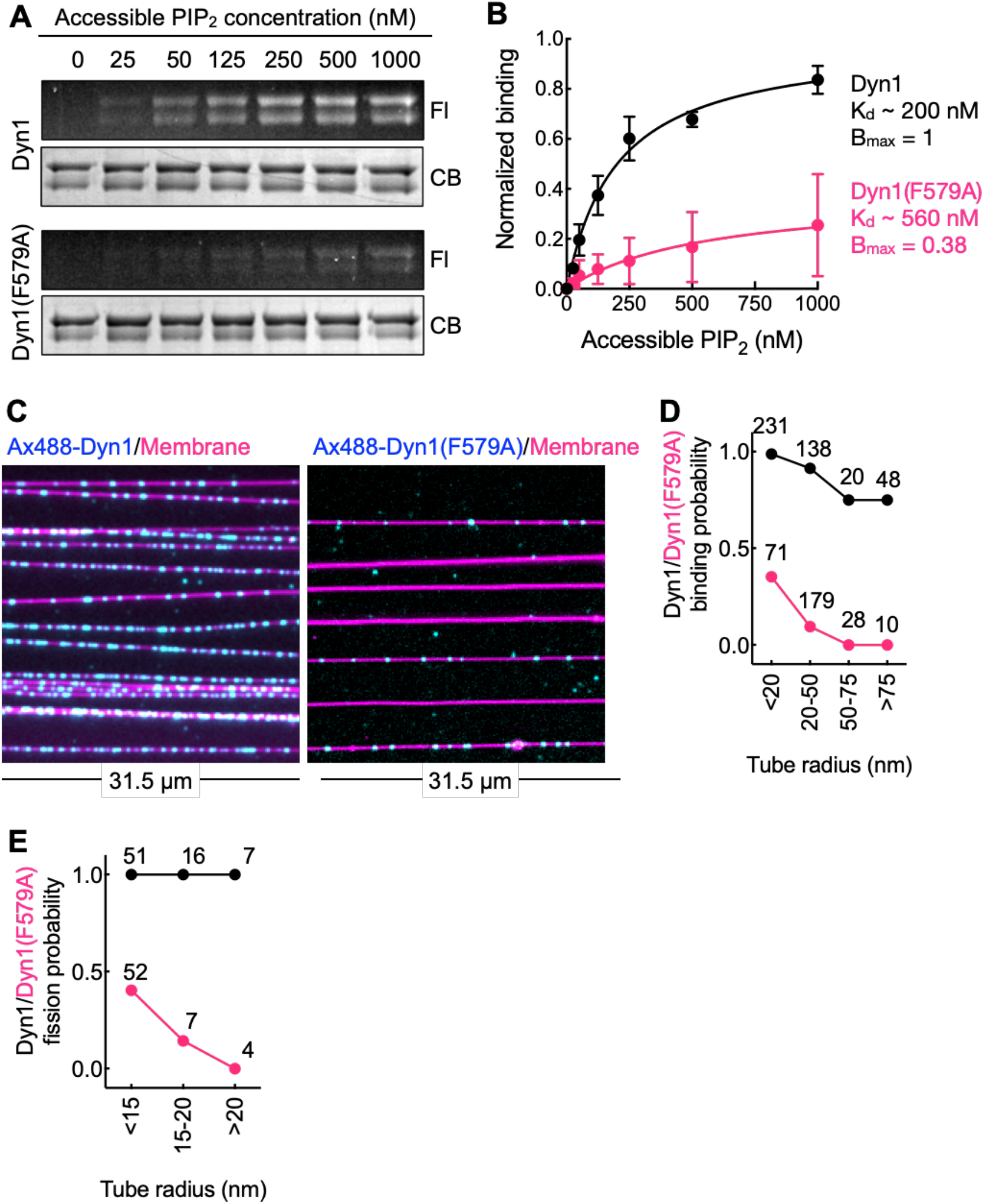
Analyzing the VL4 mutant on membranes with physiological lipid composition. (A) Results from a representative PLiMAP experiment showing in-gel fluorescence (Fl) and Coomassie brilliant blue (CB) staining of Dyn1 and Dyn1(F579A) on 1 mol% PI(4,5)P_2_ and 15 mol% DOPS liposomes. (B) Quantitation and fits of PLiMAP data with Dyn1 and Dyn1(F579A) to a one-site binding isotherm. Data represent the mean ± SD of three independent experiments. (C) Representative micrographs showing the distribution of Alexa488-labeled fluorescent Dyn1 and Dyn1(F579A) on SMrTs. (D) Quantitation of binding probability of Dyn1 and Dyn1(F579A) as a function of tube radius. The number of tubes analyzed for each size range is indicated in the plot. (E) Quantitation of fission probability with Dyn1 and Dyn1(F579A) as a function of tube radius. The number of tubes analyzed for each size range is indicated in the plot.

The above-described assays are quite minimal in the sense that they rely solely on the dynamin PHD’s ability to bind lipids for membrane recruitment. In cells, dynamin relies on multivalent interactions with lipids and endocytic proteins. The PHD binds PI(4,5)P_2_ while the PRD binds a host of SH3 domains in endocytic proteins, which together facilitate dynamin’s recruitment to the membrane. To recreate such multivalent interactions, we tried recruiting dynamin binding partner proteins on SMrTs. Our attempts with dyn1 partners like ampiphysin1 and endophilin were unsuccessful because they showed negligeable binding to SMrTs containing 1 mol% PI(4,5)P_2_ and 15 mol% DOPS. We then turned to Amphiphysin2 or BIN1 (bridging-interactor 1), specifically isoform 8 which contains a positively charged PI stretch that binds PI(4,5)P_2_ with high affinity, and an SH3 domain that interacts with dynamin and also participates in CME as a dynamin binding partner (Taylor et al., 2011; Ramjaun and McPherson, 1998; Picas et al., 2014; Hohendahl et al., 2016).

Flowing BIN1-GFP onto SMrTs containing 1 mol% PI(4,5)P_2_ and 15 mol% DOPS caused it to readily bind nanotubes. BIN1-GFP appeared uniformly distributed on some nanotubes (see box with dotted line and associated fluorescence profiles in Fig. 3A) while on others, it was organized as discrete foci (see box with solid line and associated fluorescence profiles in Fig. 3A). These foci coincided with lower membrane fluorescence, implying constriction of the underlying tube. BIN1 is widely known for its ability to tubulate planar membranes (Picas et al., 2014). But a tendency for BIN1 to organize into membrane active scaffolds on tubes has not been reported. We therefore investigated this phenomenon. The coefficient of variation (COV) of BIN1-GFP fluorescence along the length of the tube reports on the non-uniformity in BIN1 distribution. The COV showed a rise with an increase in tube radius (Fig. 3B). Conversely, BIN1-GFP membrane density, which is the average BIN1-GFP fluorescence divided by the average membrane fluorescence, showed a decline with an increase in tube radius (Fig. 3C). Together, these results indicate that an increase in tube size converts BIN1 organization from a long and continuous scaffold into small and discrete units, likely because of limiting protein density on the membrane. Indeed, raising the PI(4,5)P_2_ levels to 5 mol% caused BIN1 to show a more uniform distribution (data not shown). Correlating the tube radius under BIN1 scaffolds to the starting tube radius clearly reveals that constriction activity of BIN1. Thus, tubes of ∼10 nm starting radius retain their dimension while tubes of ∼30 nm radius get constricted to ∼12 nm (Fig. 3D). These estimates agree well with the limiting value of 14 nm reported from cryoEM measurements of large vesicles tubulated by BIN1 (Adam et al., 2015). Based on these attributes, BIN1-coated membrane nanotubes should represent an ideal template to interrogate dynamin functions in a more native context. Scaffolds represent sites of high local density of SH3 domains and their ability to constrict tubes to a narrow range of 8-14 nm radius would be expected to rescue the acute membrane curvature dependence seen for the VL4 mutant in membrane binding and fission.

**Fig. 3.**
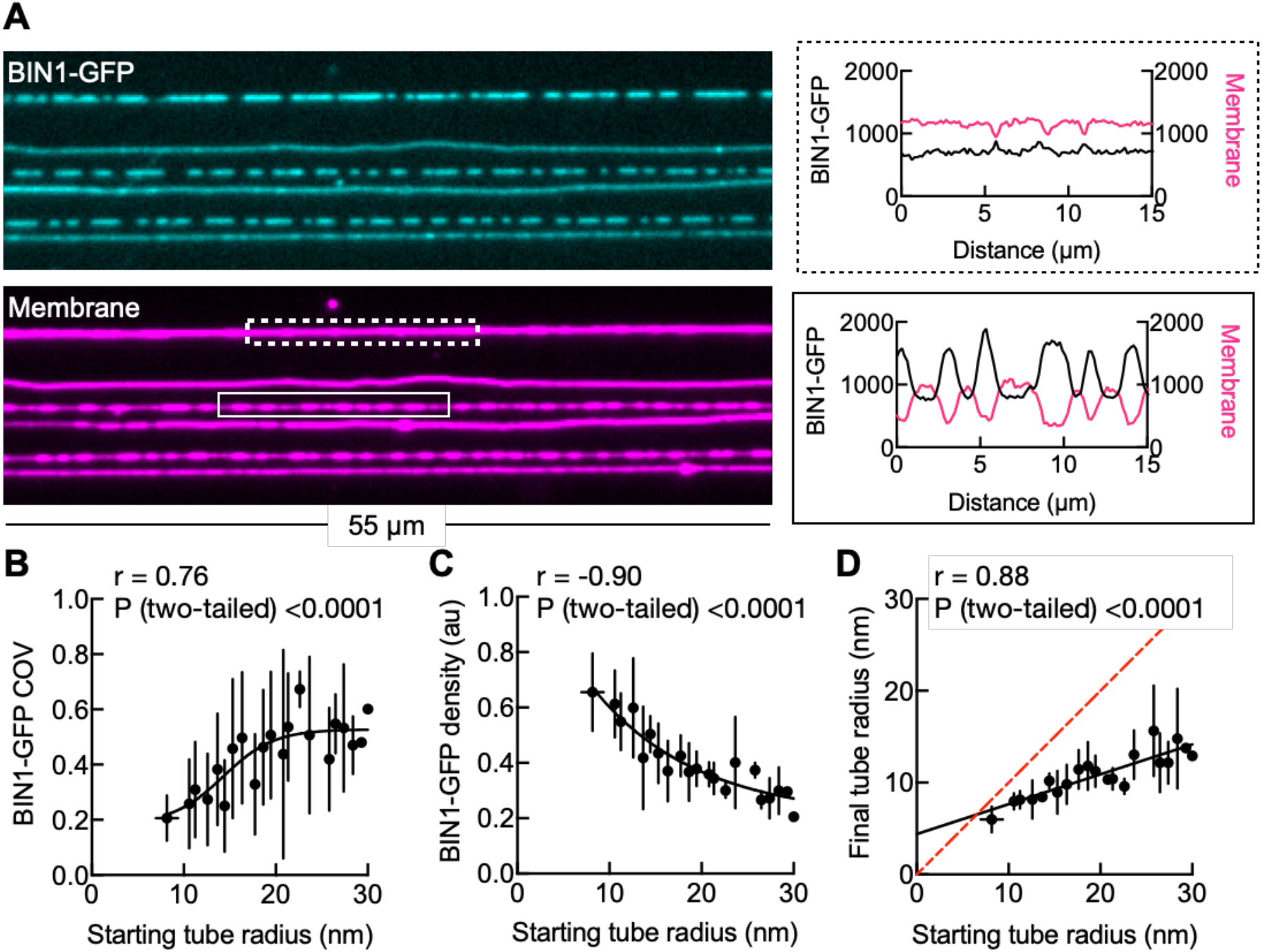
Analyzing BIN1 organization on membrane nanotubes. (A) A representative micrograph showing the distribution of BIN1-GFP on SMrTs. (B) Coefficient of variation (COV) of BIN1-GFP fluorescence as a function of tube radius. (C) BIN1-GFP density as a function of tube radius. (D) Final tube radius under BIN scaffolds as a function of the starting tube radius. The red dotted line represents a scenario without tube constriction. Data in (B-D) represent the mean ± SD with the starting tube radius estimates binned to their integral values.

To confirm if dynamin binds BIN1 via SH3-PRD interactions on membrane nanotubes, especially because of the partial C-terminal truncation of the PRD seen in our recombinant dynamin preparations (see Methods), we first prepared BIN1-GFP-coated membrane nanotubes and flowed in fluorescent Dyn1 and Dyn1(F579A). Dynamin fluorescence coincided with BIN1-GFP fluorescence implying efficient binding (Fig. 4A). Thus, the partial C-terminal truncation of the PRD appears not to significantly impact dynamin’s ability to engage with BIN1’s SH3 domain. We then assayed for dynamin functions in presence of GTP on BIN1-coated membrane nanotubes. For these experiments, we monitored only the membrane fluorescence channel to improve temporal resolution and because our results confirm that BIN1 is either distributed uniformly on narrow membrane tubes or is localized as discrete scaffolds, each of which coincide with dimmer tube fluorescence (Fig. 3). Flowing Dyn1 with GTP on BIN1-coated membrane nanotubes showed robust fission. Fission was apparent on nanotubes that were uniformly coated with BIN1 as well as on those displaying localized BIN1-dependent constrictions (Fig. 4B). On tubes showing localized BIN1-dependent constrictions, time-lapse imaging revealed that fission took place within the BIN1 scaffold, which is evident from the dimmer constricted region undergoing splitting (Supplementary Fig. 2). Remarkably, flowing Dyn1(F579A) with GTP showed no fission of BIN1-coated membrane nanotubes (Fig. 4B). Indeed, correlating fission probability to the starting tube size revealed this mutant to be fission-defective across a range of tube sizes (Fig. 4C).

**Fig. 4.**
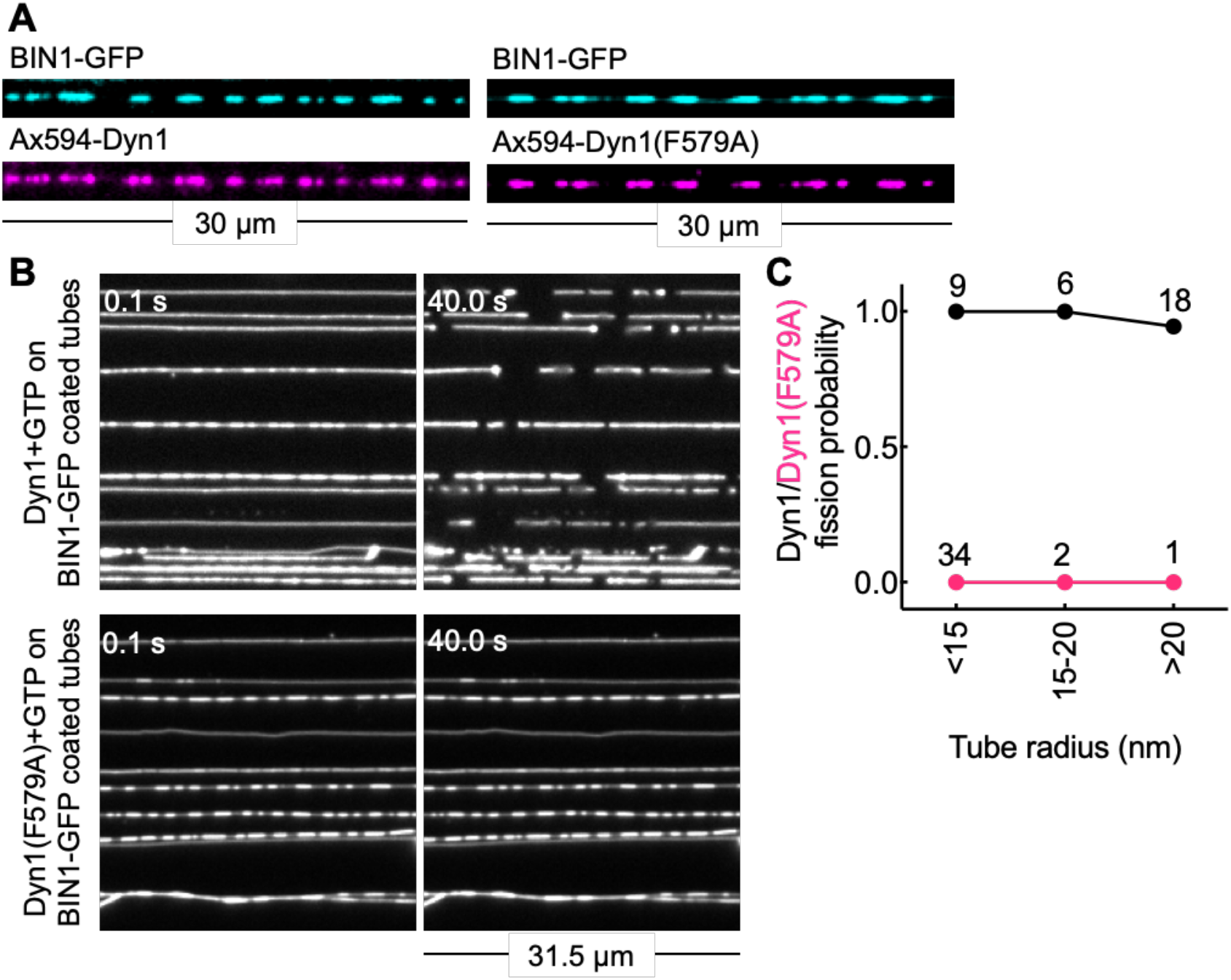
Analyzing the VL4 mutant on BIN1-coated membrane nanotubes. (A) Representative fluorescence micrographs of SMrTs showing localization of Alexa594-labeled Dyn1 and Dyn1(F579A) on BIN1 scaffolds. (B) Representative fluorescence micrographs showing the effect of addition of Dyn1 and Dyn1(F579A) with GTP to BIN1-coated membrane nanotubes. (F) Quantitation of fission probability with Dyn1 and Dyn1(F579A) as a function of starting tube radius. The number of tubes analyzed for each size range is indicated in the plot.

Finally, we tested the VL4 mutant in a cellular assay for dynamin functions. Dynamin’s ability to sustain CME has been monitored using clathrin-dependent uptake of transferrin. During late stages of clathrin-mediated endocytosis, dynamin is recruited to necks of clathrin-coated pits to catalyze fission leading to the release of clathrin-coated vesicles (Taylor et al., 2011). Expression of GTPase-, self-assembly and membrane binding-defective dynamin mutants causes the arrest of transferrin uptake by dominant-negative inhibition of native dynamin function (Damke et al., 1994; Ramachandran et al., 2009; Marks et al., 2001). This experimental paradigm is also relevant to understand potential CMT-linked mutations since these are autosomal dominant. We tested the importance of VL4 by monitoring transferrin uptake in cells overexpressing Dyn1(F579A)-GFP and Dyn1-GFP. These assays were carried out in dynamin2 KO HeLa cells to specifically test dyamin1 functions. Cells overexpressing Dyn1(F579A)-GFP showed a significant reduction in transferrin uptake compared to cells expressing Dyn1-GFP. This is apparent from the intense perinuclear fluorescence of transferrin seen in control and Dyn1-GFP expressing cells but absent in Dyn1(F579A)-GFP expressing cells (Fig. 5A). Quantification of these results revealed a significant 2.6-fold reduction in transferrin uptake in cells expressing Dyn1(F579A)-GFP compared to that seen upon overexpression of Dyn1-GFP, implying a strong dominant-negative inhibition of native dynamin functions. These results corroborate data from in vitro assays showing a defect in membrane fission and signify the importance of VL4 for dynamin functions.

**Fig. 5.**
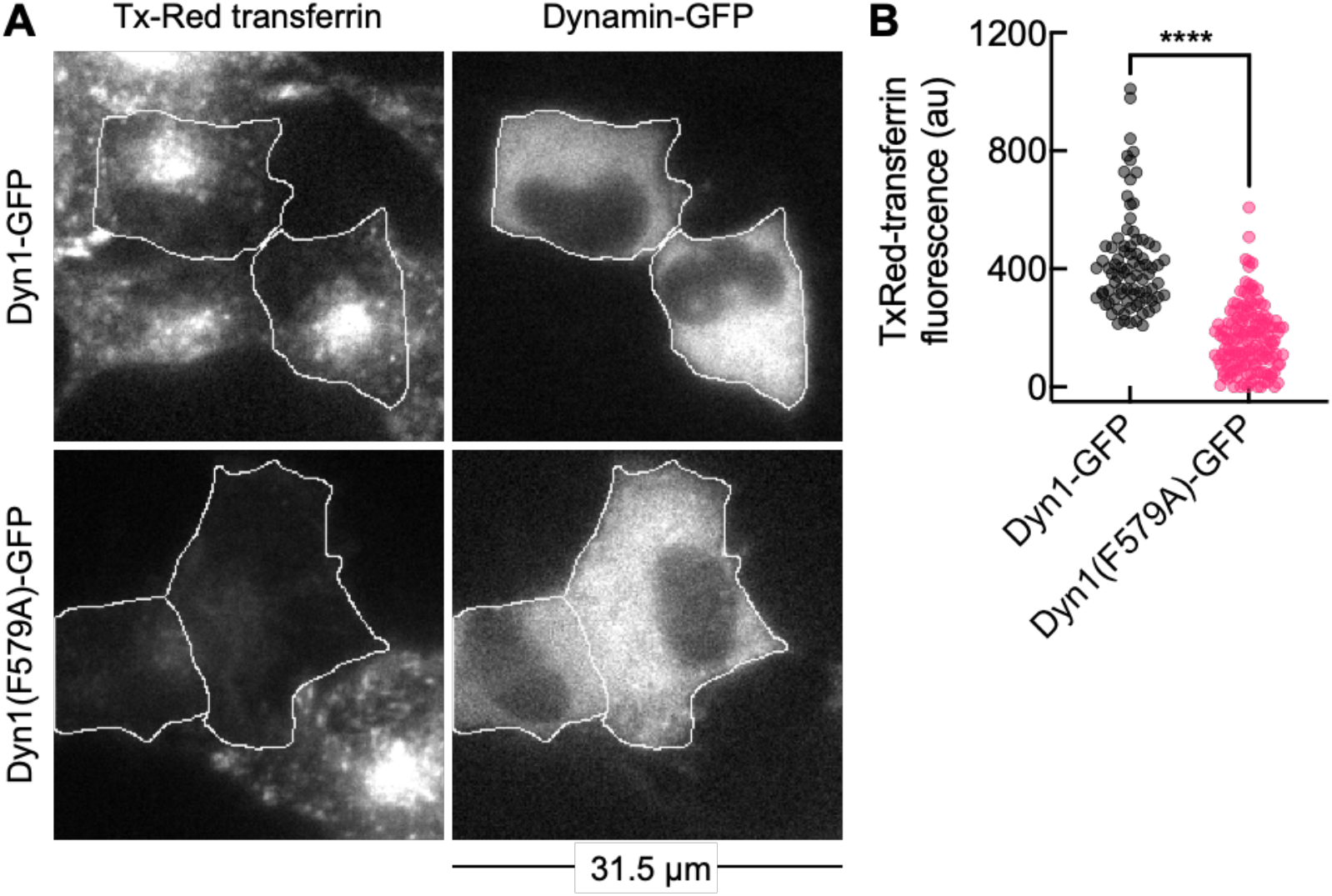
The VL4 mutant is defective in clathrin-mediated endocytosis. (A) Representative micrographs showing uptake of fluorescent transferrin in dynamin2 KO HeLa cells expressing Dyn1-GFP and Dyn1(F579A)-GFP. (B) Quantitation of florescent transferrin uptake in Dyn1-GFP expressing cells (n=84) and Dyn1(F579A)-GFP expressing cells (n=148) in two independent experiments. Significance was estimated using Mann-Whitney test where **** denotes P<0.0001.

## Discussion

Recent MD simulations show that the isolated PHD primarily binds the membrane through residues in the VL1 and VL4 loops. While the essential role of VL1 has been well established, the present work is the first to experimentally address the importance of VL4 to dynamin functions. Guided by simulations showing that the F579 residue in VL4 approaches closest to the membrane, we tested the F579A mutation for functions. In bulk liposome-based assays, Dyn1(F579A) displays significant defects in binding to membranes containing physiologically relevant levels of PI(4,5)P_2_. Extending such analysis to a spatially resolved microscopic assay on membrane nanotubes, we find that the F579A mutation renders Dyn1 binding more sensitive to membrane curvature. Interestingly, simulations of the F579A mutant in the isolated PHD showed no apparent defect in membrane binding. This discrepancy is likely because membrane dissociation would be favored in the full-length protein than with an isolated PHD such that a subtle reduction in the hydrophobicity of VL4 could change membrane partitioning properties more significantly for the full-length protein than the isolated PHD.

Dynamin self-assembles into helical scaffolds and its binding is inherently favored on membranes of high curvature (Roux et al., 2010). But dynamin scaffolds constrict tubes thereby extending the range of curvatures that can support binding and self-assembly. Our results indicate that the F579 residue in VL4 contributes to this process since when mutated, dynamin shows a steep decline in binding with a decrease in membrane curvature. However, on tubes of a size range that can recruit Dyn1(F579A), addition of the mutant with GTP caused fission indicating that the mutant is partially defective in function. We expected that the defect in membrane binding and fission in Dyn1(F579A) would be rescued by involving SH3-PRD interactions in addition to lipid-based interactions. Instead, we find that Dyn1(F579A) functions are further affected with its fission capacity completely abolished on BIN1 scaffolds. BIN1 scaffolds are expected to facilitate dynamin functions. They represent sites with a high local concentration of SH3 domains and, based on results presented here, also constrict tubes to a fission-compliant size. Previous reports have indicated that a stoichiometric excess of BAR domain proteins such as endophilin and ampiphysin can inhibit dynamin functions, likely by forming mixed scaffolds that prevent dynamin’s G domain interactions that are necessary for stimulated GTPase activity (Hohendahl et al., 2017; Takei et al., 1999). Our assays are designed to allow BIN1 to first form scaffolds and then recruit dynamin. Washing off excess BIN1 before dynamin addition could have ensured that dynamin self-assembles within the BIN1 scaffold to cause fission. Such sequential addition more closely mimics the cellular scenario wherein endocytic proteins arrive before dynamin for fission of clathrin-coated pits (Daumke et al., 2014; Taylor et al., 2011). Dyn1(F579A) bound BIN1 scaffolds but surprisingly fails at fission suggesting that the PHD interactions with the membrane are important. The PHD actively facilitates membrane fission. This is evident from our previous work showing that replacing the native PHD-PI(4,5)P_2_ interaction with a generic 6xHis-chelator lipid interaction or with a polylysine-PS interaction in dynamin supports fission but with slower kinetics and with the presence of long-lived highly constricted tubular intermediates (Dar and Pucadyil, 2017). This is likely because VL insertion has been shown to facilitate nonbilayer-like arrangements of membrane lipids and lower the membrane bending energy (Fuhrmans and Müller, 2015; Baratam et al., 2021; Shnyrova et al., 2013). Despite its localization to the BIN1 scaffold, the lack of fission seen with Dyn1(F579A) could reflect an inability for the PHD to actively engage with the membrane. For tubes of similar sizes, Dyn1(F579A) causes fission without BIN1 but is inhibited with BIN1. This effect could have emerged because BIN1 scaffolds compete with dynamin for PI(4,5)P_2_ binding. In such a scenario, Dyn1(F579A) would be affected more than Dyn1 because of its lower affinity for PI(4,5)P_2_. This could emphasize a general principle wherein endocytic partner proteins act in a mutually competing manner wherein protein-based interactions facilitate dynamin’s recruitment while lipid-based interactions compete with dynamin’s engagement with the membrane. Consequently, subtle defects in membrane binding could become amplified within the local microenvironment of the BIN1 scaffold thus rendering Dyn1(F579A) defective in fission and CME.

The PHD is a hotspot for mutations that cause several genetic disorders. Many of the mutations reside in regions that are highly conserved across all isoforms of dynamin. Our work extends these observations by characterizing functions of the VL4 in dynamin. The missense mutation M580T in VL4 that causes the CMT neuropathy could arise from a decrease in VL4 hydrophobicity (Haberlová et al., 2011). A testament to the importance of this loop is apparent from the fact that a more subtle F579A mutation than M580T causes a dramatic effect on dynamin function. Our results therefore emphasize the importance of this novel VL4 membrane anchor in dynamin functions and reveal how finely tuned lipid-protein interaction have evolved to facilitate dynamin functions in membrane fission and CME.

## Acknowledgements

HK and SB thank IISER Pune for graduate fellowships. We thank Mike Ryan (Monash University) for the dynamin2 KO HeLa cells. TJP thanks the HHMI for funding support. We thank the Pucadyil Lab members and Anand Srivastava (Indian Institute of Science) for comments on the manuscript.

## Materials and methods

### Constructs and plasmids

For bacterial expression, human dynamin1 was cloned in pET15b with N-terminal 6xHis and C-terminal StrepII tags. The F579A mutation was introduced using PCR. Human BIN1-EGFP (isoform 8) (Addgene plasmid #27305) was cloned in pET15b with N-terminal 6xHis and C-terminal StrepII tags. For mammalian cell expression, human dynamin1 and the F579A mutant in pET15B was cloned with a C-terminal GFP fusion into pcDNA3.0. All clones were confirmed by sequencing.

### Protein purification and fluorescent labeling

Proteins were expressed in BL21(DE3) cells grown in autoinduction medium at 18°C for 36 h. Bacterial cells were pelleted and stored at -40 °C. The frozen bacterial pellet was thawed in 20 mM HEPES pH 7.4, 500 mM NaCl with a protease inhibitor cocktail tablet (Roche) and lysed by sonication in an ice-water bath. Lysate were spun at 30,000 g for 20 min and the supernatant was incubated with HisPur™ Cobalt Resin (Thermo Fisher Scientific). The resin was washed with 20 mM HEPES pH 7.4, 500 mM NaCl and bound protein was eluted with 20 mM HEPES pH 7.4, 500 mM NaCl, 100 mM EDTA. The elution was loaded onto a StrepTrap HP column (GE Lifesciences), washed with 20 mM HEPES pH 7.4, 150 mM NaCl and eluted with the same buffer containing 2.5 mM desthiobiotin. Despite this two-step purification procedure, dynamin elutes with a partial C-terminal truncation likely because it exists as a tetramer (Dar and Pucadyil, 2017). Proteins were spun at 100,000 g to remove aggregates before use in assays. Protein concentration was estimated from UV absorbance at 280 nm using the molar extinction coefficient predicted by the Expasy ProtParam tool. Dynamin was labeled with Alexa488 C5-maleimide or Alexa594 C5-maleimide (Invitrogen) using a 5-fold excess of the reactive dye in 20 mM HEPES pH 7.4, 150 mM NaCl buffer. The reaction was quenched with excess DTT and the unreacted dye was removed by extensive dialysis.

### Liposome preparation

1,2-dioleoyl-sn-glycero-3-phosphocholine (DOPC), 1,2-dioleoyl-sn-glycero-3-phospho-L-serine (sodium salt), (DOPS), and 1,2-dioleoyl-sn-glycero-3-phospho-(1’-myo-inositol-4’,5’-bisphosphate) (ammonium salt) (18:1 PI(4,5)P_2_) were from Avanti Polar Lipids. The UV-activable, diazirine-containing fluorescent lipid probe (BODIPY-diazirine PE) was prepared as described earlier (Jose and Pucadyil, 2020; Jose et al., 2020). Lipids were aliquoted at desired ratios in a glass tube and dried under high vacuum for 30 min to a thin film. Deionized water was added to the dried lipids to achieve a final concentration of 1 mM. Lipids were hydrated at 50 °C for 30 min, vortexed vigorously and extruded through 100 nm pore-size polycarbonate filters (Whatman).

### GTPase assays

For GTPase assays, dynamin (0.1 μM) was incubated with DOPS liposomes (10 μM) in 20 mM HEPES pH 7.4, 150 mM KCl buffer containing 1 mM GTP (Jena Bioscience) and 1 mM MgCl_2_ and the released inorganic phosphate was measured using a malachite green-based colorimetric assay (Baykov et al., 1988; Dar and Pucadyil, 2017).

### Proximity-based labeling of membrane-associated proteins (PLiMAP)

PLiMAP was carried out as described earlier (Jose and Pucadyil, 2020; Jose et al., 2020). Briefly, liposomes containing 1 mol% of BODIPY-diazirine PE were incubated with dynamin at a 100:1 lipid:protein molar ratio in a final volume of 30 μl. The reaction was incubated in the dark at room temperature for 30 min and exposed to 365 nm UV light (UVP crosslinker CL-1000L) at an intensity of 200 mJ cm^-2^ for 1 min. The reaction was mixed with sample buffer, boiled and resolved using SDS-PAGE. Gels were first imaged for BODIPY fluorescence on an iBright1500 (Thermo Fischer Scientific) and later fixed and stained with Coomassie Brilliant Blue. Binding data was fitted to a one-site binding isotherm using Graphpad Prism.

### Supported Membrane Templates (SMrT), fluorescence imaging and image analysis

Supported membrane templates (SMrT) were prepared as described earlier (Dar et al., 2017). Briefly, lipids were aliquoted at desired ratios to a final concentration of 1 mM in chloroform. The lipid mixes also contained the fluorescent lipid *p*Texas-Red DHPE (Thermo Fisher Scientific) at 1 mol% concentration. 2 μl of the lipid mix was spread with a glass syringe on a PEGylated glass coverslip, dried, and assembled inside an FCS2 flow chamber (Bioptechs). The chamber was filled with 20 mM HEPES pH 7.4, 150 mM NaCl and flowed at high rates to form SMrTs. For binding, 0.3 μM dynamin was flowed onto SMrTs in 20 mM HEPES pH 7.4, 150 mM NaCl and incubated for 10 mins. Templates were imaged after washing off excess dynamin. For fission, SMrTs were pre-equilibrated with an oxygen scavenger cocktail in 20 mM HEPES pH 7.4, 150 mM NaCl and time-lapse images were acquired while flowing in 0.3 μM dynamin mixed with 1 mM GTP (Jena Bioscience) and 1 mM MgCl_2_ in 20 mM HEPES pH 7.4, 150 mM NaCl. For experiments involving BIN1, 0.2 μM of BIN1-GFP was flowed onto SMrTs, incubated for 10 mins and unbound protein was washed off before flowing in dynamin with or without GTP. Tubes radii were estimated as described earlier (Dar et al., 2017). Imaging was carried out through a 100x, 1.4 NA oil-immersion objective on an Olympus IX83 inverted microscope connected to an LED light source (CoolLED) and an Evolve 512 EMCCD camera (Photometrics). Image acquisition was controlled by µManager and analyzed using Fiji (Schindelin et al., 2012).

### Cell culture, live cell imaging and transferrin uptake assay

Dynamin2 KO HeLa cells have been reported earlier (Kamerkar et al., 2018). Cells were cultured in complete DMEM with 10% FBS and 1% penicillin-streptomycin (HiMedia) and maintained at 37 °C in humidified 5% CO_2_. Cells were transfected with dynamin using Lipofectamine 2000 (Thermo Fisher Scientific). Transfected cells were trypsinized and plated on 40 mm glass coverslips (Bioptechs). Transferrin uptake experiments were performed between 24-48 hrs post-transfection. Cells were serum-starved for 2 hrs in serum-free DMEM and the coverslips were assembled in an FCS2 chamber maintained at 37 °C. The media was exchanged for HEPES-buffered Hank’s balanced salt solution and cells were fed with 50 μg.ml^-1^ of Texas Red-labelled transferrin (Invitrogen) and incubated for 10 mins. Excess transferrin was washed off and the cells were imaged through 60x, 1.4 NA oil-immersion objective on an Olympus IX83 inverted microscope attached with a stable LED light source (CoolLED) and an Evolve 512 EMCCD camera (Photometrics), using appropriate excitation wavelength and emission filters for Texas Red and GFP. Image acquisition was controlled by Micro-Manager and rendered using Fiji (Schindelin et al. 2012).

---

**Supplementary Fig. 1.**
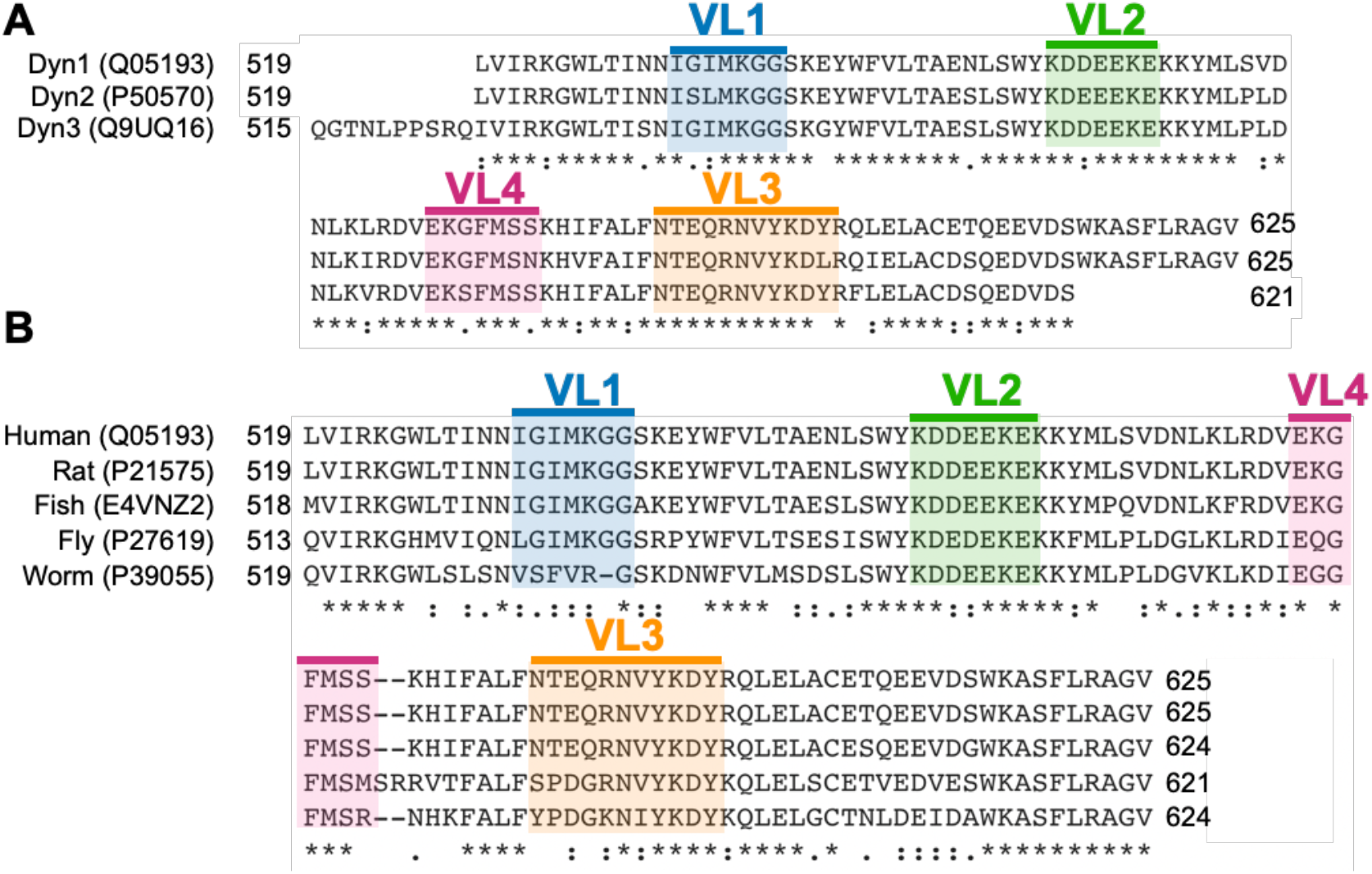
Sequence alignment of the dynamin PHD. Sequence alignment of the PHD in dynamin isoforms in human (A) and dynamin PHD orthologues from different species (B) with the VL1-4 marked.

**Supplementary Fig. 2.**
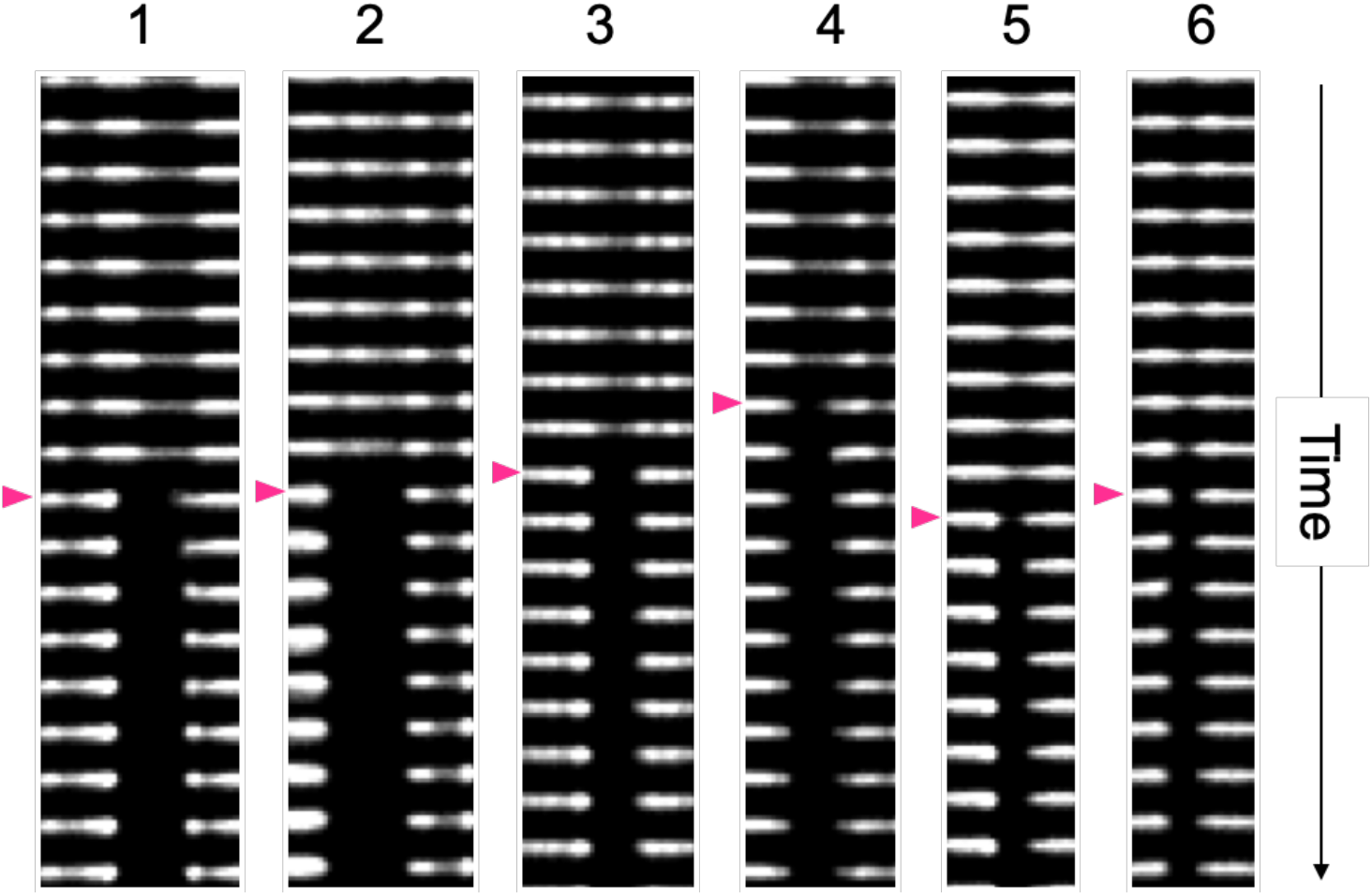
Dynamin-catalyzed fission of BIN1-coated membrane nanotubes. The panel shows 6 independent fission events showing that the site of fission is localized within the BIN1 scaffold, which is apparent from the dimmer constricted region on the nanotube undergoing splitting. Red arrowhead marks the time frame when fission is observed.

## Notes

### Competing Interest Statement

The authors have declared no competing interest.

